# *nkx3.2* mutant zebrafish accommodate jaw joint loss through a phenocopy of the head shapes of Paleozoic jawless fish

**DOI:** 10.1101/776351

**Authors:** Tetsuto Miyashita, Pranidhi Baddam, Joanna Smeeton, A. Phil Oel, Natasha Natarajan, Brogan Gordon, A. Richard Palmer, J. Gage Crump, Daniel Graf, W. Ted Allison

## Abstract

The vertebrate jaw is a versatile feeding apparatus that facilitated explosive diversification. To function, it requires a joint between the upper and lower jaws, so jaw joint defects — such as osteoarthritis or even ankylosis — are often highly disruptive and difficult to study. To describe consequences of jaw-joint dysfunction, we engineered two independent null alleles of a single jaw-joint marker gene, *nkx3.2*, in zebrafish. These mutations caused zebrafish to become functionally jawless via fusion of the upper and lower jaw cartilages (ankylosis). Despite lacking jaw joints, *nkx3.2* mutants survive to adulthood and accommodate this defect by: a) remodeling their skulls; and b) altering their behavior from suction feeding to ram feeding. As a result of remodeling, *nkx3.2* mutants developed superficial similarities to the skull shapes observed in two lineages of ancient jawless vertebrates (anaspids and furcacaudiid thelodonts), including: a fixed open gape, reduced snout, and enlarged branchial region. However, no homology exists in individual skull elements between these taxa, and most of the modified elements in the mutant zebrafish occur outside known expression domains of *nkx3.2*. Therefore, we interpret the adult *nkx3.2* phenotype not as a reversal to an ancestral state, but as convergence due to similar functional requirement of feeding without moveable jaws. This remarkable convergence strongly suggests that jaw movements themselves dramatically influence the development of jawed vertebrate skulls, which implies that functionally viable skull morphologies are finite, with or without functional jaws. Because *nkx3.2* null zebrafish display prominent joint ankylosis, drastically modified skull shape, and altered feeding behaviors, these mutants provide a unique model with which to investigate mechanisms of skeletal remodeling and joint diseases.

## INTRODUCTION

The jaw is a functionally versatile innovation that facilitated explosive diversification of gnathostomes (a clade containing jawed vertebrates), but its basic structure is surprisingly simple and highly conserved (Miyashita, 2016). A jaw consists of ‘a hinge and caps’: upper and lower skeletal levers hinged at a jaw joint (Depew and Simpson, 2006). As the joint enables biting motions, its origin is considered the final step in the evolutionary assembly of the vertebrate jaw (Cerny et al., 2010; Kuratani, 2012; Miyashita, 2016). Across jawed vertebrates, the presumptive jaw joint is marked by the expression of *nkx3.2*, an NK2 class homeobox gene (a.k.a. *bapx*), at the midheight of the embryonic mandibular arch (Gillis et al., 2013; Lukas and Olsson, 2018a; Miller et al., 2003; Tucker et al., 2004). Chondrogenesis dorsal to this expression domain gives rise to a palatoquadrate (upper jaw), whereas chondrogenesis ventral to it forms Meckel’s cartilage (lower jaw) (Medeiros and Crump, 2012). This basic pattern remains conserved among jawed vertebrates, but later development varies. Marginal bones arise intramembranously around the often endo-/peri-chondrally ossified jaw cartilages except in chondrichthyans (sharks, rays, and skates) (Hall, 2015). In mammals, the jaw joint instead forms between two such intramembranous bones (temporal and dentary), whereas the proximal jaw joint becomes the malleus-incus interface that is, in mice, no longer affected by *Nkx3.2* knockout (Tucker et al., 2004). Despite these variations after pharyngeal chondrogenesis, no gnathostome lineage secondarily lost functional jaws.

By studying functional jaw loss in our new mutant zebrafish, we asked whether — and how — jaw functions affect vertebrate skull shape during development. Clinically documented agnathia in humans typically accompanies severe congenital disorders such as holoprosencephaly and otocephaly, but jaw loss is clearly a secondary effect and not a cause in these cases (Bixler et al., 1985; Brown and Marsh, 1990; Gekas et al., 2010; Schiffer et al., 2002). Instances of temporomandibular joint ankylosis (stiffening due to bone fusion) may result from trauma or infection, or may be congenital (Adekeye, 1983; Chidzonga, 1999; Manganello-Souza and Mariani, 2003). If untreated, the ankylosis can lead to the ‘bird face’ deformity (El-Sheikh et al., 1996). However, these cases do not fully document the effects of functional jaw loss. In mammalian models, various jaw/skull deformations have been induced by surgical resection, detachment, or repositioning of the jaw muscles and/or bones (Bayram et al., 2010; Gomes et al., 2012; Horowitz and Shapiro, 1955; Lifshitz, 1976; Miyazaki et al., 2016; Rodrigues et al., 2009; Sarnat, 1970; Sarnat and Muchnic, 1971; Toledo et al., 2014). These manipulations occurred well after formation of the jaw skeleton and muscles, and the jaws remained partially functional because of unilateral operations or non-comprehensive disruption. The defects and deformities reported in these studies imply: a) jaw movements are potentially an important factor in shaping the skulls; and b) any allele disrupting jaw movements would be generally maladaptive. Nevertheless, these implications are difficult to explore without an accessible experimental model.

To fill this gap, we engineered two distinct null alleles of *nkx3.2* in zebrafish. Previously, transient knockdown of *nkx3.2* during early development (using morpholinos) had shown fusion of the nascent jaw cartilages in both zebrafish and frogs (Lukas and Olsson, 2018a; Miller et al., 2003). We confirmed in zebrafish that the mutants reproduce this phenotype. Surprisingly, mutant zebrafish are viable — despite loss of the jaw joint — and grow through to adulthood. Functionally jawless as a result, *nkx3.2*^-/-^ zebrafish dramatically alter skull shape late in ontogeny to facilitate feeding, with the mouth fixed open, the snout reduced, and the branchial region expanded. This open-mouth phenotype, previously unknown in zebrafish or any other jawed vertebrates, also occurred in two extinct lineages of 400-million-plus year-old jawless vertebrates, anaspids and thelodonts. Even though they share no homology in individual facial bones, *nkx3.2*^-/-^ fish accommodate loss of a functional jaw by converging onto these ancient, distantly related agnathan head shapes. Thus, *nkx3.2* mutant zebrafish provide a unique model for both skeletal remodeling and joint diseases such as osteoarthritis, and to reevaluate evolutionary implications of phenocopies in general.

## MATERIALS AND METHODS

### Animal Ethics

Zebrafish maintenance and experiments were approved as protocol number AUP00000077 by the Animal Care and Use Committee: Biosciences at the University of Alberta as dictated by the Canadian Council on Animal Care. Other zebrafish work was approved by the University of Southern California Institutional Animal Care and Use Committee.

### Animal husbandry

Embryos were incubated at 28 °C, and treated with 0.003% PTU (1-phenyl-2-thiourea) in 10% Hank’s saline starting at 24 hpf. Larvae were introduced to the nursery at 1 week to 10 dpf. Genomic DNA was extracted from clipped fins of 3-5 dpf larvae or from adults. Preserved tissues, embryos, larvae, and adults were all fixed in 4% PFA, and stored in 100% EtOH or MeOH at −20 °C except for adults (preserved in 70% EtOH at 4°C).

### Molecular genetics

Nkx3.2 protein is a transcription factor with a homeobox DNA binding domain that is 100% conserved in amino acid sequence among zebrafish, mouse, and human homologs. In zebrafish, a single *nkx3.2* gene is apparent in the genome, and its homology to mammalian *NKX3.2* is strongly supported by gene synteny: e.g. the neighbor genes flanking *nkx3.2* on zebrafish Chromosome 14 (*wdr1* and *bod1l1*) are positioned coordinately in mouse, human and spotted gar.

Two disparate regions of the gene were targeted by CRISPR guide RNA (gRNA) or TALENs, producing two disparate null alleles that produced similar phenotypes (Fig. 1A). One allele (ua5011) was engineered with CRISPR/Cas9 (Gagnon et al., 2014) targeted at the beginning of the homeodomain, and it harbors a 20 bp deletion resulting in a frameshift (Fig. 1A; Data Supplement 1). The disrupted translation of codons is predicted to abrogate production of the critical homeobox domain, and instead produce random amino acids. This is predicted to produce a non-functional Nkx3.2 and a null allele. A disparate allele (el802) was generated using TALENs (Barske et al., 2016) targeted at the start of the gene. This produced a stably inherited gene with 20 bp deletion, removing the translation start codon (Fig. 1A). The allele *nkx3.2^el802^* is predicted to not produce Nkx3.2 protein. Morphologically, these two alleles are not readily distinguishable from each other (see Results).

**Fig. 1.**
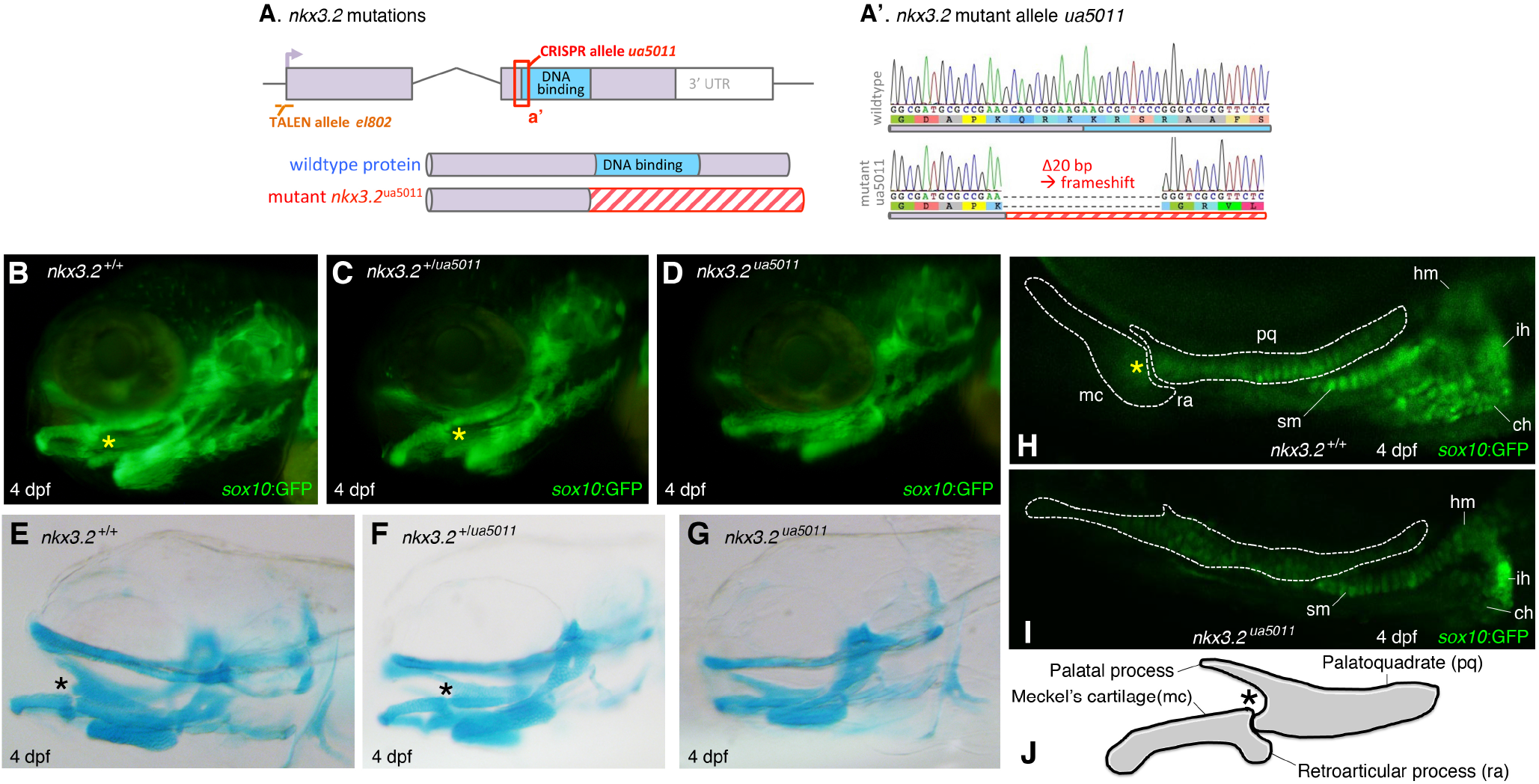
Null alleles of *nkx3.2* in zebrafish result in the jaw joint ankylosis. (**A**) Mutations engineered with CRISPR/Cas9 or TALEN in the zebrafish gene *nkx3.2* that encodes the homeobox transcription factor protein Nkx3.2 (a.k.a. Bapx1). <u>Top:</u> The gene *nkx3.2* contains two exons (purple), the second of which encodes the homeobox domain (DNA binding domain, blue). The diagram is approximately to scale (exon 1 = 310 bp), except the 3’ untranslated region (3’UTR). Allele *nkx3.2^el802^*was generated with TALEN technology that deleted 20 bp from the start of the gene (orange dashed line), eliminating the start codon. Allele *nkx3.2^ua5011^* was engineered with CRISPR/Cas9 to generate a 20 bp deletion (red box) and frameshift that is predicted to eliminate the homeodomain. <u>Bottom:</u> Schematic of predicted protein following CRISPR mutagenesis. In the allele *nkx3.2^ua5011^*, the frameshift (20 bp deletion) is predicted to disrupt the translation and abrogate production of the critical homeobox domain. This is predicted to produce random amino acids (red hatching). (**A’**) Sequencing results from allele *nkx3.2^ua5011^* (lower) compared to a wildtype zebrafish (upper). At 4 dpf, the chondrocrania are compared by *sox10*:eGFP expression in chondrocytes (**B**–**D**, **H**, **I**) or by alcian blue staining of cartilages (**E**–**G**) between age-matched specimens. (**B**, **E**) The chondrocranial morphology of wildtypes (AB background; *sox10*:eGFP) at 4 dpf in left lateral view, in *sox10*:eGFP expression within chondrocytes (B) and alcian blue staining of cartilages (E). (**C**, **F**) The chondrocranial morphology of *nkx3.2* heterozygous mutants (*nkx3.2*^+/*ua5011*^; *sox10*:eGFP) at 4 dpf in left lateral view, in *sox10*:eGFP expression within chondrocytes (C) and alcian blue staining of cartilages (F). (**D**, **G**) The chondrocranial morphology of *nkx3.2* homozygous mutants (*nkx3.2^ua5011^*; *sox10*:eGFP) at 4 dpf in left lateral view, in *sox10*:eGFP expression within chondrocytes (D) and alcian blue staining of cartilages (G). (**H**, **I**) Comparison of jaw morphology between age-matched wildtype (AB background; *sox10*:eGFP) (H) and *nkx3.2* homozygous mutants (*nkx3.2^ua5011^*; *sox10*:eGFP) (I) at 4 dpf in left lateral view, using *sox10*:eGFP expression within chondrocytes. White broken lines delineate the mandibular cartilages. (**J**) Schematic drawing of the mandibular cartilages in zebrafish at 4 dpf in left lateral view, showing wildtype morphology (see panel E). **Abbreviations: asterisk (*)**, jaw joint; **aa**, anguloarticular; **ar**, articular; **bh**, basihyal; **ch**, ceratohyal; **d**, dentary; **dpf**, days post fertilization; **hm**, hyomandibula; **ih**, interhyal; **j**, jugal; **ke**, kinethmoid; **lp**, lower lip; **mc**, Meckel’s cartilage; **m**, maxilla; **n**, nostril or aperture of nasohypophyseal system; **o**, operculum; **pm**, premaxilla; **po**, preopercular; **pq**, palatoquadrate; **q**, quadrate; **ra**, retroarticular process; **sm**, symplectic; **up**, upper lip.

### CRISPR

To generate *nkx3.2<u>^ua5011^</u>*, we designed sgRNA to five different targets near or within the *nkx3.2* homeodomain, which were all injected:

GGCGGCCATCTGACGTCGCT
GGCTGACGCCAGCAGATCGG
AAGCAGCGGAAGAAGCGCTC
GAGCGCTTCTTCCGCTGCTT
GGCCGCGTTCTCCCACGCGC

These targets were selected using the web resource CHOPCHOP (Labun et al., 2016; Montague et al., 2014). Following the protocol developed by Gagnon and colleagues (2014), two different oligonucleotides were ordered: one containing a target sequence led by the SP6 promoter (ATTTAGGTGACACTATA) and followed by the overlapping region (GTTTTAGAGCTAGAAATAGCAAG) of the reverse oligonucleotide; and the reverse containing the constant, Cas9-binding domain of sgRNA. These oliogonucleotides were annealed after 5-minute incubation at 95°C, through graded cooling (−2°C s^-1^ to 85°C; −0.1°C s^-1^ to 25°C), and filled in for the non-overlapping regions using T4 DNA polymerase (NEB: M0203S). To synthesize sgRNAs using these templates, MegaScriptTM SP6 Transcription Kit (Ambion: AM1330) was used. The RNAs were precipitated in ammonium acetate solution, suspended in UltraPure^TM^ H_2_O, and stored in 2-3 µl aliquots at −80 °C. For injection, sgRNA(s) were diluted to 400-600 ng µl^-1^, with 1 µl mixed with 1 µl aliquot of Cas9 nuclease from *Streptococcus pyogenes* (NEB: #M0646) at 1 µg ml^-1^. This solution was mixed with 3 µl of 0.2M KCl, 0.2 % phenol red, and ddH_2_O. The final injection volume per embryo was approximately 5 nl, with 400-600 pg sgRNA and 1 ng Cas9 nuclease. For control, GFP 5’GA was used at the stage P_0_ when ubi:switch/RH+AB was crossed, which can be phenotyped by reduction of ubiquitous GFP in the progenies with dsRed expression in heart.

We generated *nkx3.*2^ua5011^ against AB background. Cas 9 and sgRNAs targeted for *nkx3.2* were coinjected with sgRNA that disrupts GFP GA5’ (CTCGGGCTGAAGCTCGGCG), at stages between fertilization and first cleavage, to fertilized eggs collected from the crossing of ua3140 ubi:switch/AB+RH and the background AB line. At 3 dpf, injected larvae were sorted for reduced expression of ubiquitous GFP and the presence of dsRed fluorescence in heart. These larvae provided the P_0_ population. Sequencing of genomic DNA extracted from fin clips of the P_0_ adults identified a female with a 20 bp deletion to the homeodomain-coding region of *nkx3.2* (ua5011). The P_0_ female carrying this mutation was crossed to the *sox10*:GFP transgenic line, and progenies were sorted at 3 dpf for the presence of *sox10*:GFP expression and the absence of the two markers (ubiquitous GFP expression and red fluorescent heart). ua5011 heterozygotes were identified by both sequencing of extracted gDNA and Restriction Fragment Length Polymorphism (RFLP) analysis using one XmaI (NEB: R01805) restriction site within the deleted region (primers for genotyping: 5’– GGACGAGACGGATCAGGAATC–3’; 5’–CACTCGGCGTGTTCGGTAAA–3’). These F_1_ heterozygotes were incrossed for F_2_ embryos, which were genotyped by RFLP analysis and phenotyped at 4 dpf by identifying *sox10*:GFP-positive chondrocytes and staining cartilages using alcian blue. Homozygotes were reared with a strictly small-grained diet to the adult stage. In this study, *nkx3.2*^ua5011/ua5011^ represent F_2_ generation derived from the P_0_ mutant female and a wildtype male (AB; *sox10*:GFP), whereas comparative wildtype (AB; *sox10*:GFP) come from incrossing of half-siblings of the P_0_ male.

### Tissue preparation and histology

One- and two-month-old *nkx3.2^+/+^* and *nkx3.2*^-/-^ zebrafish were fixed in 4% paraformaldehyde for 24hrs. Zebrafish were eviscerated prior to decalcification with 0.5M ethylenediaminetetraacetic acid (EDTA) solution for 4 weeks. Samples were dehydrated post decalcification in a series of graded ethanol and embedded in paraffin. Tissue blocks were embedded in a sagittal orientation and sections were cut at 7µm using a 820 Spencer microtome. Hematoxylin and eosin staining was performed on zebrafish sections by firstly placing sections in an oven at 60 °C for 10 min. The deparaffinized sections were rehydrated using xylene and graded ethanol (100%, 95%, 70%), followed by staining with hematoxylin and eosin. The slides were then dehydrated and mounted using Permount.

### Skeletal preparation

Alcian blue staining of cartilages partly followed the protocol provided by Michael Shapiro (University of Utah). Specimens fixed in 4% PFA were rinsed with ddH_2_O and transferred to 70% EtOH. Once equilibrated, larvae were immersed in alcian blue solution (0.167 mg/ml alcian blue; 15% acetic acid; 70% EtOH), rinsed through EtOH/ddH_2_O series, and washed in a saturated sodium borate solution. Specimens were immersed in trypsin solution (0.125% trypsin; 30% sodium borate) overnight, washed in 1% KOH solution, bleached in 0.15% H_2_O_2_ 0.1% KOH, 25% glycerol solution, immersed through a 1% KOH/glycerol graded series into 100% glycerol for storage. Specimens older than 21 dpf were immersed in 0.005% alizarin red solution in 1% KOH overnight, after the first 1% KOH wash and before bleaching in 0.15% H_2_O_2_ 0.1% KOH.

### Filming

Wildtype (AB; *sox10*:GFP) and *nkx3.2*^ua5011/ua5011^ were filmed at 2 months post fertilization to record feeding behavior (Movie S1). A fish was placed in a 1.4L tank with dark background and given brine shrimp larvae. Feeding was filmed using Canon EOS 760D at 30 frames s^-1^ in dimensions 1280 × 720 pixels. The films were cropped and assembled using iMovie (ver. 10.1.9, © 2001-2018 Apple Inc.) and slowed to 1/10 original speed.

### Imaging

#### Micro-computed tomography (µCT)

Two-month-old zebrafish were scanned using MILabs µCT scanner. Scans were reconstructed at a voxel size of 25µm. Images were analyzed using AVIZO 3-dimensional software (Milabs, Utrecht, Netherlands). 2-Dimensional images used for linear and geometric morphometrics were obtained from AVIZO.

#### Microscopy

Fluorescent images were acquired on a Zeiss Axio Observer.Z1 with LSM 700 confocal microscope via ZEN 2010 software (version 6.0, Carl Zeiss MicroImaging, Oberkochen). Brightfield imaging of stained preps was performed on a Leica MZ16F dissection microscope (Concord ON, Canada) with 12.8 megapixel digital camera (DP72, Olympus; Richmond Hill ON, Canada).

### Morphometrics

#### Rationales for morphometric comparison

The purpose of our quantitative comparison is to test phenotypic similarities and differences qualitatively identified in *nkx3.2*^-/-^ mutants with respect to wildtype zebrafish and anaspids (and thelodonts for gape angles). On the one hand, the skulls of *nkx3.2*^-/-^ mutants clearly depart from wildtype morphologically at adult stage (Fig. 2). To describe this morphological departure in greater details, we will present comparison of skeletal growth between *nkx3.2*^-/-^ and wildtype elsewhere. On the other hand, it is difficult to assess observed similarities between *nkx3.2*^-/-^ phenotype and the general head configuration in anaspids. Zebrafish and anaspids are distant to each other phylogenetically: the former is nested deep within, in the ascending order, cypriniforms, teleosts, neopterygians, actinopterygians, osteichthyans, and gnathostomes, whereas anaspids represent either a stem gnathostome or even a stem cyclostome lineage (Donoghue et al., 2000; Janvier, 2007, 1996; Keating and Donoghue, 2016; Miyashita et al., 2019). The dermatocranium of a zebrafish is macromeric, although that of an anaspid is largely micromeric with scales of acellular bone (Blom et al., 2001). There is no morphological correspondence in individual elements of the skull roof between zebrafish and anaspids. The parabranchial cavity is closed by the operculum in zebrafish, whereas each branchial pouch had its own outlet in the series of external pores in anaspids (Blom et al., 2001). A single gene mutation in *nkx3.2* did not reverse these, and other morphological differences accumulated after the last common ancestor of zebrafish and anaspids. For morphometric comparison, we chose metric traits that can be identified in both zebrafish and anaspids.

**Fig. 2.**
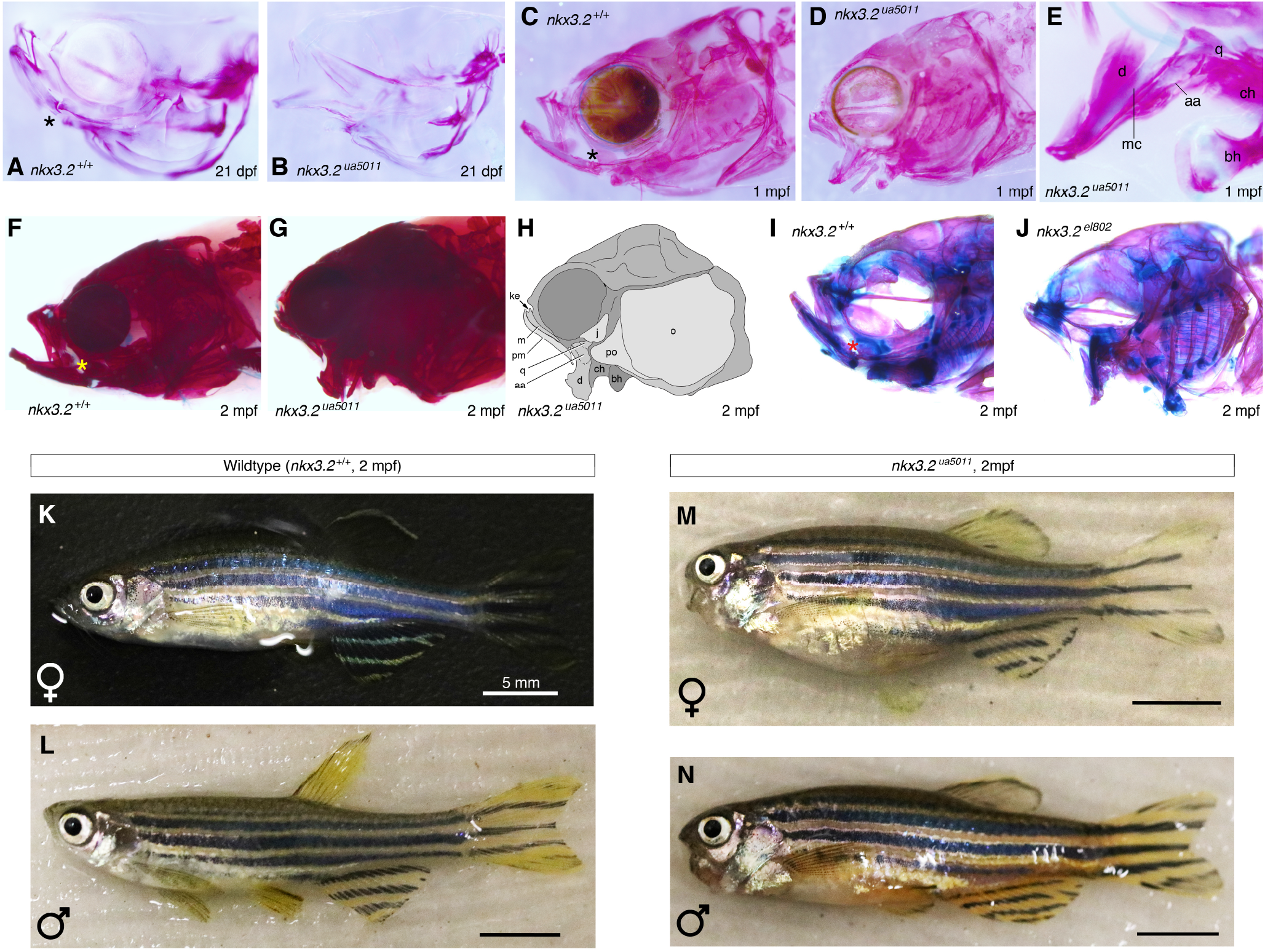
Ontogeny of *nkx3.2^-/-^* zebrafish documents drastic remodeling of the skull. (**A, B**) Comparison of the skull morphology between age-matched wildtype (AB strain) (A) and *nkx3.2* homozygous mutant (*nkx3.2^ua5011^*) (B) at 21 dpf in right lateral view (inverted for consistency with other panels), using alcian blue and alizarin red staining. (**C**, **D**) Comparison of skull morphology between age-matched wildtype (AB) (C) and *nkx3.2* homozygous mutant (*nkx3.2^ua5011^*) (D) at 1mpf in left lateral view, using alcian blue and alizarin red staining. (**E**) Detailed morphology of jaw skeleton in *nkx3.2* homozygous mutant (*nkx3.2^ua5011^*) at 1 mpf in right lateral view (inverted for consistency with other panels), using alcian blue and alizarin red staining. (**F**, **G**) Comparison of skull morphology between age-matched wildtype (AB) (F) and *nkx3.2* homozygous mutant (*nkx3.2^ua5011^*) (G) at 2 mpf in left lateral view, using alcian blue and alizarin red staining. (**H**) Interpretive drawing of the specimen in panel g. (**I**, **J**) Comparison of skull morphology between age-matched wildtype (AB) (I) and *nkx3.2* homozygous mutant (*nkx3.2^el802^*) (J) at 2 mpf in left lateral view, using alcian blue and alizarin red staining. This second independent null allele of *nkx3.2* confirms the phenotypes reported herein. (**K**–**N**) Comparison of overall morphology at 2 mpf among age-matched wildtype (AB) female (K) and male (L) and *nkx3.2* homozygous mutant (*nkx3.2^el802^*) female (M) and male (N) in left lateral view. Scale bars = 5 mm. Abbreviations follow Fig. 1.

#### Linear morphometrics

The fixed open gape is similar between *nkx3.2*^-/-^ mutants and anaspids. We compared this trait by taking the angle between supporting skeletal elements of the upper and lower lips in both taxa. In zebrafish, the angle was taken by extrapolating the axis of the premaxilla until it meets the axis of the dentary. At stages younger than the onset of dermal ossification (4 and 14 dpfs), the angle was measured between the axes of the palatal process (palatoquadrate) and Meckel’s cartilage. In anaspids, the upper and lower lips are demarcated by a series of plates or relatively larger scales, which allowed delineation of the gape angle (measured at where the extrapolated upper and lower lip margins meet). The angle was measured similarly in thelodonts, except that the lip margins were identified along the series of small marginal scales. Gape angle exhibits a roughly normal distribution within each of the age class of both *nkx3.2*^-/-^ and wildtype zebrafish and among anaspids (see Data Supplement 2). In addition to gape angles, we measured lengths of skulls and lower jaws in zebrafish, and orbit diameter in anaspids and thelodonts. Original measurements are available in Data Supplement 2.

In *nkx3.2*^-/-^ zebrafish, the gape angle increases progressively with age, and thus with increasing body size. Anaspids and thelodonts overall have much greater range of body size than zebrafish. Unlike *nkx3.2*^-/-^ zebrafish, however, the gape angle appears to vary independently of body size in anaspids. No correlation exists between gape angle and orbit diameter in anaspids, regardless of whether among those specimens falling in the size range of zebrafish or across the entire clade (*r* = −0.114; *P* = 0.306). Although eye size is generally negatively allometric in vertebrates (Howland et al., 2004), the orbit diameter is one reliable, structurally intact metric trait in this clade, because most specimens are not preserved in entire body length, and because other reference measurements (e.g., body height) are affected by taphonomic deformation or simply not preserved in most specimens (Blom et al., 2001; Blom and Märss, 2010; Janvier, 1996; Sansom et al., 2010). In thelodonts, the relationship remains to be tested between body size and gape angles because of small sample size (*n* = 5).

#### Geometric morphometics

Eight landmarks were assigned to both zebrafish (2 mpf) and anaspid samples for geometric morphometric comparison. These landmarks capture general configuration of the heads (1: anterior tip of upper lip; 2: junction between upper and lower lips; 3: anterior tip of lower lip; 4: nostril, or nasohypophyseal opening; 5: anterior extremity of orbit; 6: posterior extremity of orbit; 7: trunk-head boundary at dorsal outline; 8: ventral point of hypobranchial region). These landmarks describe structures homologous across vertebrates (landmarks 4, 5, 6) or geometrically determined positions comparable across vertebrates. They are free of morphological discontinuity between anaspids and gnathostomes (none of the landmarks represents anaspid- or gnathostome-specific morphology). TpsDig (Rohlf, 2018) was used to place landmarks on two-dimensional images of anaspid and zebrafish. The digitized file was entered into MorphoJ software (Klingenberg, 2011) and all images were aligned using the anterior and posterior extremity of the orbits. These landmark data were transformed using the procrustes superimposition method (Rohlf, 1999), and the resulting coordinates were compared using Principal Component Analysis (PCA). In PCA, PC scores from anapsid, *nkx3.2^+/+^* and *nkx3.2*^-/-^ zebrafish were grouped by equal frequency ellipses with a P value of 0.95.

#### Rationales for selection of comparative taxa in geometric morphometrics

From the pool of nearly a thousand catalogued specimens of anaspids, we selected a total of 70 specimens that show lateral compression during the fossilization process, with the least taphonomic artifact, to reflect lateral view of the heads. Furcacaudiid thelodonts were excluded from geometric morphometrics because only a handful of exceptionally preserved specimens are available. The sample size is small for this latter group (*n* < 5), and all such specimens were collected from a single locality, making it difficult to identify (and thus control for) taphonomic artifacts. In addition, landmarks cannot be assigned confidently in this group. The nasohypophyseal opening cannot be located precisely because of the micromeric nature of the integument on the dorsal side of the head (Wilson and Caldwell, 1998, 1993). The transition from head to trunk is ambiguous along the dorsal outline because there is no apparent change in morphology of the scales (Wilson and Caldwell, 1998, 1993).

There are many other lineages of jawless vertebrates, including living cyclostomes, heterostracans, thelodonts, galeaspids, pituriaspids, and osteostracans, in the order of nested hierarchy toward the crown-group gnathostomes (Janvier, 2007, 1996). Living cyclostomes are difficult to compare as they have an anguilliform profile and lack ossified skeletons, or cartilages unambiguously comparable to the jaw/lip elements of the zebrafish skull (Miyashita, 2016, 2012; Miyashita et al., 2019). Extinct jawless vertebrates generally have a depressiform profile (Janvier, 1996). Therefore, direct shape comparison is difficult with the compressiform zebrafish and anaspids. Among non-depressiform jawless stem gnathostomes, the lips are typically preserved poorly. Furcacaudiid thelodonts have a lateromedially compressed body profile, and have the lip morphology consistent with that of anaspids (Wilson and Caldwell, 1998, 1993). These similarities suggest that depressed lower lips are a general condition along the gnathostome stem, as reconstructed conventionally across the stem group.

## RESULTS

### Jaw joint is ankylosed in *nkx3.2* null alleles

At 4 days post fertilization (dpf), *nkx3.2*^-/-^ zebrafish replicated the *nkx3.2* morpholino knockdown phenotype: the absence of a jaw joint (Miller et al., 2003). The palatoquadrate and Meckel’s cartilage fused together, and the retroarticular process was absent (Fig. 1D, G, I). Apart from joint ankylosis, no marked differences were apparent in overall morphology or survival rates between *nkx3.2*^-/-^ mutants and wildtype (some minor difference in skull size and lower jaw proportions are discussed in the next section). Heterozygotes were morphologically indistinguishable from wildtype (Fig. 1C, F), and the *nkx3.2* alleles displayed recessive Mendelian inheritance (F_2_ genotypes followed Mendelian ratio; heterozygotes developed wildtype morphology). Remarkably, these functionally jawless homozygous mutants survived beyond early larval stages.

### Functionally jawless *nkx3.2*^-/-^ zebrafish modify skull shapes late in ontogeny

Contrary to the maladaptive nature of jaw dysfunctions in general, *nkx3.2*^-/-^ zebrafish continued to grow without a jaw joint. Marked phenotypic differences against wildtype began to emerge between the 2^nd^ and 3^rd^ weeks post fertilization (Fig. 2). The lower jaw became downturned in *nkx3.2*^-/-^ fish resulting in a rigidly fixed open mouth, whereas both upper and lower jaws were upturned in wildtype zebrafish (Fig. 2A, B). This timing coincides with the onset of ossification and the period of active feeding in normal juveniles (Cubbage and Mabee, 1996; Kimmel et al., 1995). Although the palatoquadrate and Meckel’s cartilage remained fused in *nkx3.2*^-/-^ mutants, skeletal staining reveals that the upper and lower jaw elements ossified independently of each other — still without a ball-and-socket joint structure (Fig. 3A, B). All skull elements in *nkx3.2*^-/-^ mutants ossified without apparent delay.

**Fig. 3.**
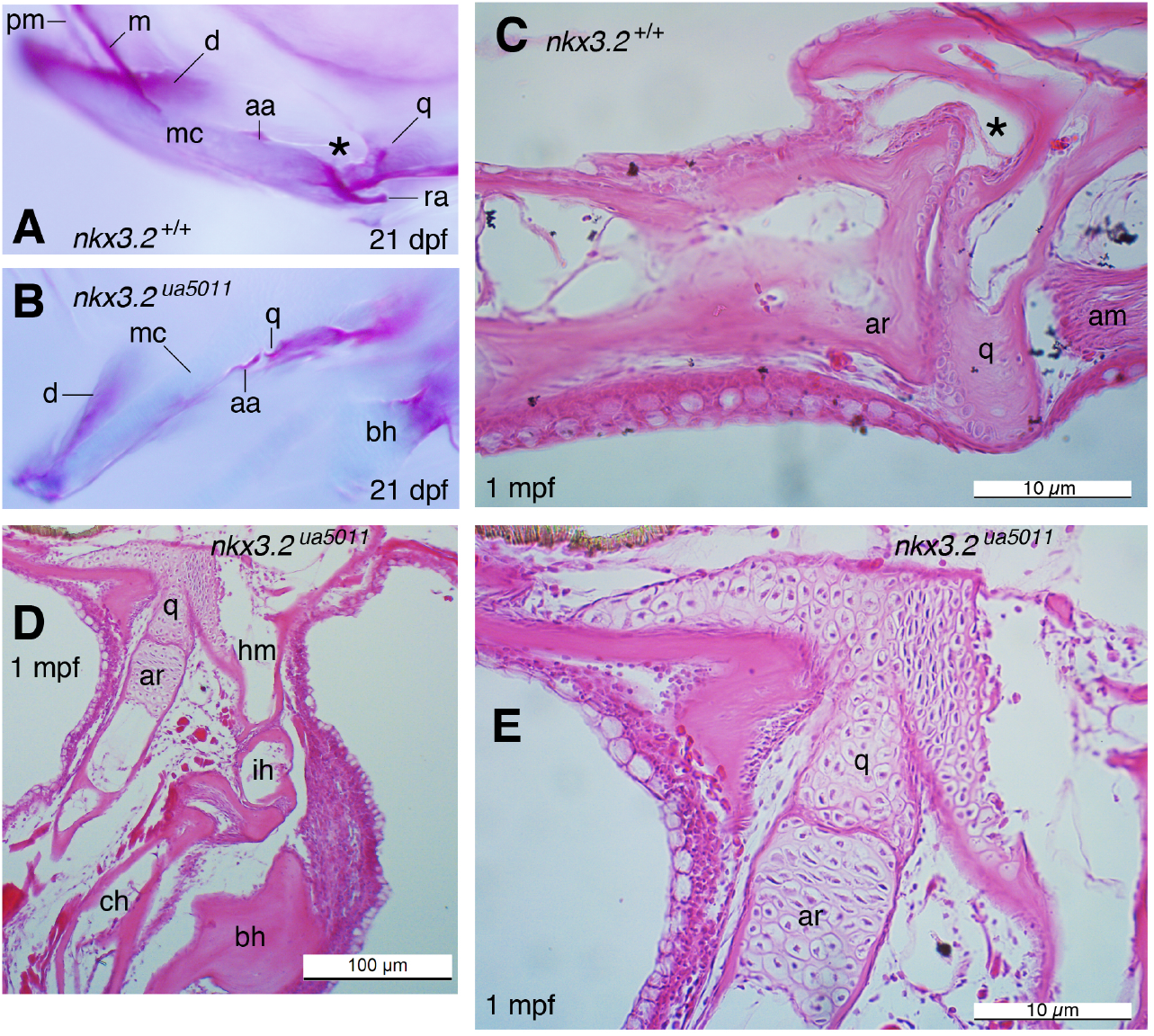
Detailed morphology of jaw joint in wildtype, or the joint-less interface between the upper and lower jaws in *nkx3.2*^-/-^ mutants. (**A, B**) Detailed morphology of junction between upper and lower jaws in age-matched wildtype (A) and *nkx3.2* homozygous mutant (*nkx3.2^ua5011^*) (B) at 21 dpf in left lateral view, using alcian blue and alizarin red staining. (**C**, **D**) Sagittal section of junction between upper and lower jaws in age-matched wildtype (AB) (C) and *nkx3.2* homozygous mutant (*nkx3.2^ua5011^*) (D) at 1 mpf, stained with eosin and hematoxylin. (**E**) Detailed view of section in panel D.

The lower jaws were increasingly turned downward in *nkx3.2*^-/-^ by the end of the first month, resulting in a greater gape (Fig. 2D, E). The jaw joint was still absent in *nkx3.2*^-/-^ mutants: a sheet of perichondrium lies between the ossifying quadrate and articular — so the two bones remained distinct elements — but this interface had none of those essential components of synovial diarthrosis (Fig. 3D, E). Many other osteological differences emerged by this stage. Normally, the premaxilla and the maxilla swing forward to sit nearly vertical and are hinged by the kinethmoid for suction feeding (Hernandez, 2000; Hernandez et al., 2007) (Fig. 2C). In *nkx3.2*^-/-^ mutants, however, the premaxilla and the maxilla became oriented posteroventrally and abutted against the anterior margin of the orbit (Fig. 2D). The kinethmoid was reduced into a fused bony process, unlike a rod-like hinge element in wildtype (Hernandez et al., 2007). The downturned lower jaws of the *nkx3.2*^-/-^ mutants were relatively shorter than the normal lower jaws of wildtypes. The basihyal protruded anteroventrally as much as the lower jaw, implying that the muscle connecting those two elements (m. intermandibularis posterior) (Schilling and Kimmel, 1997) may be responsible for the lower jaw orientation. As a result of these modifications, *nkx3.2*^-/-^ mutants had a shorter snout, a fixed open gape, and a dorsoventrally tall profile.

Linear morphometrics corroborated the departure from normal morphology in *nkx3.2*^-/-^ mutants in the latter half of the first month (14 dpf onward). For absolute size, no significant difference (*P* > 0.05) in skull length emerged between wildtype and *nkx3.2*^-/-^ mutants except at 4 dpf (*t* = 2.202; *P* = 0.338) (Fig. 4B). Also at this stage, the lower jaws appear to be shorter relative to skull length in the mutants than in the wildtypes (*t* = 2.809; *P* = 0.0078) (Fig. 4C). These minor but statistically significant differences at 4 dpf may be a direct consequence of the ankylosis between palatoquadrate and Meckel’s cartilage. By the second week, however, differences between the mutants and wildtypes became non-significant in these metric traits. The lower jaw depression (gape angle > 45°) in the mutants was pronounced at 21 dpf (*t* = −11.834; *P* << 0.01) (Fig. 4A), but proportional changes to lower jaw lengths in the same mutants were only expressed in significant magnitude at 1 month (21 dpf: *t* = 1.7225; *P* = 0.091559; 30 dpf: *t* = 10.56; *P* << 0.01) (Fig. 4C). This lag between the two traits indicates that the rate of skeletal growth in the jaws followed changes in their orientation for a greater gape (and thus resulting in functional shift). Morphological variations within a cohort of *nkx3.2*^-/-^ mutants were greater at this stage than in any other, as indicated by the range of variation in orientations and relative lengths of the lower jaws (Fig. 4A, C).

**Fig. 4.**
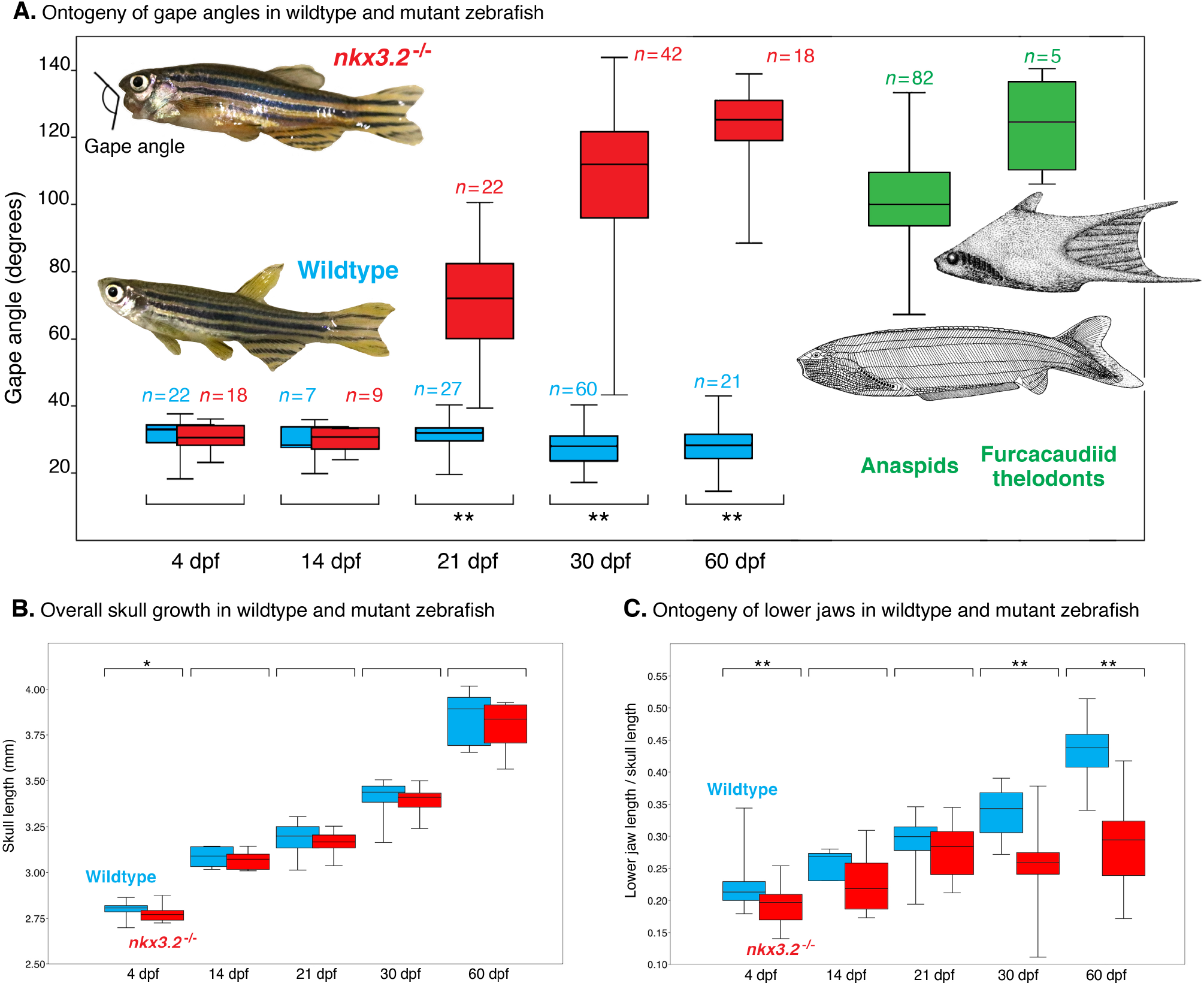
Growth of *nkx3.2*^-/-^ mutants in linear and proportional traits. (**A**) A marked departure from normal morphology (wildtype = blue) occurs in lower jaw orientations of *nkx3.2*^-/-^ zebrafish (= red) past 14 dpf, coinciding in timing with metamorphosis (onset of intramembranous ossification in the skulls and active feeding). Fixed open gapes in the mutants at 1 to 2 mpf are comparable to those of Paleozoic agnathan lineages, birkeniiform anaspids and furcacaudiid thelodonts (green). The orientations were measured here as gape, an angle between upper and lower lips at natural, resting position. **(B)** Wildtype and *nkx3.2*^-/-^ zebrafish do not differ significantly from each other in absolute sizes, except at 4 dpf. Here, skull length is selected to illustrate this general observation. (**C**) Proportional changes appear to follow shape changes in skeletal remodeling. The box plot shows phenotypic separation in relative lower jaw length between wildtype and *nkx3.2*^-/-^ zebrafish at 1 and 2 mpf, even though a significant difference developed in lower jaw orientation by 21 dpf. Values are plotted as boxes of first and third quartile, with middle line displaying mean, and whiskers communicating maximum and minimum values (*n* = sample size, same across A–C). Asterisk indicates level of statistically significant difference in means (*t*-test): *, *P* < 0.05; **, *P* < 0.01. Photographs of zebrafish are male representative specimens at 60 dpf (Fig. 2L, N). Drawings are: *Pharyngolepis oblonga* as a general representative of anaspids (after Blom et al., 2001; Kiaer, 1924); and *Furcacauda fredholmae* as a general representative of furcacaudiid thelodonts (after Wilson and Caldwell, 1993). See Data Supplement 2 for all original measurements.

### The adult *nkx3.2*^-/-^ phenotype accommodates functional jaw loss

At 2 months of age and approaching sexual maturity nearly all *nkx3.2*^-/-^ mutants had a gape angle greater than 90 degrees (Fig. 4A). These *nkx3.2*^-/-^ adults continued to be characterized by the morphological differences identified at 1 mpf. The nostrils sat between the eyes because the snout was reduced in length relative to wildtype zebrafish. The skulls appeared to be more highly ossified in *nkx3.2*^-/-^ mutants than in wildtype, where massive bones and cartilages were identified around the interface of quadrate and articular, in the lower branchial region, and in the laterally expanded operculum (Figs. 2J, G, M, N, 5B; Movie S1). All homozygous mutants showed the descriptive skeletal traits identified here. Morphologically, the phenotypic effects of the two alleles (*nkx3.2^ua5011^* and *nkx3.2^el802^*) were virtually indistinguishable from each other at respective ages (Fig. 2G, J), given variations within a strain (for examples, see variation in gape angles, relative lengths of lower jaws, or sexual dimorphism in *nkx3.2^ua5011^*: Figs.2M, N, 4A, C).

**Fig. 5.**
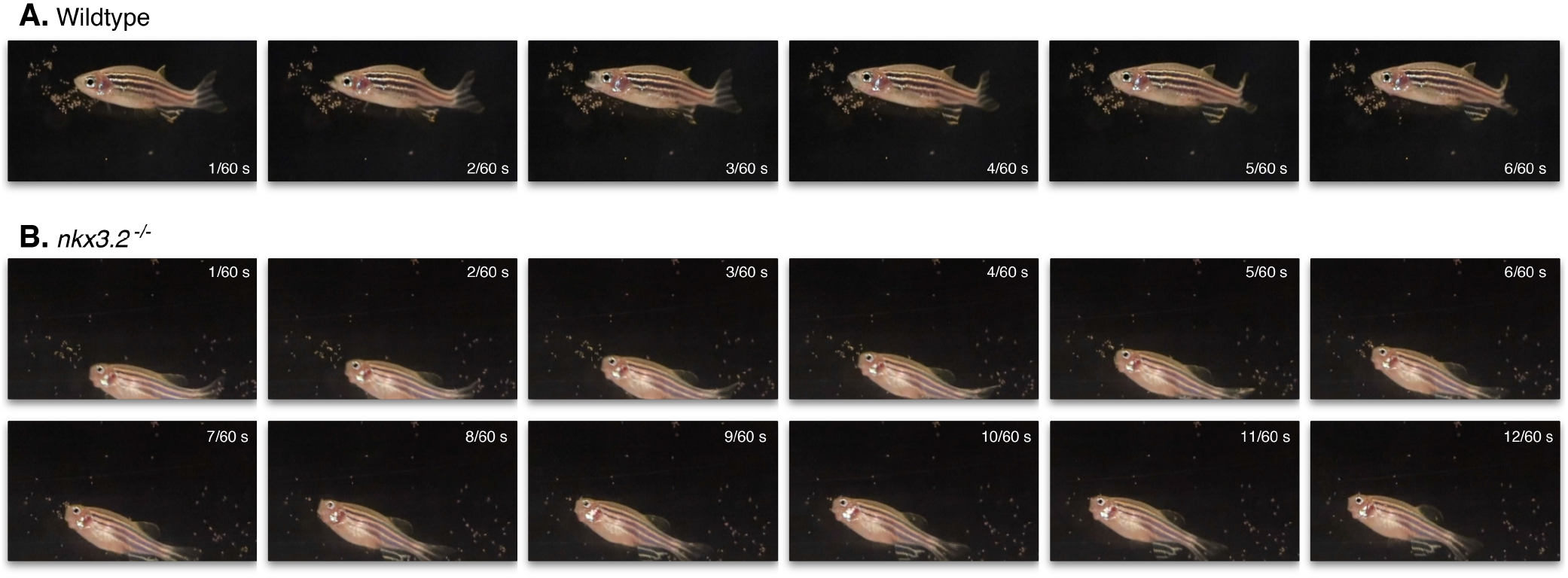
Functionally jawless *nkx3.2*^-/-^ mutants perform ram feeding. (**A**) Wildtype zebrafish (6 mpf) use suction feeding, following the general principles of suction feeding mechanics of actiopterygians: lower jaw depression, forward swing of premaxilla and maxilla, expansion of parabranchial cavity, and recoiling motions in that order. An entire cycle takes approximately 0.1 s. (**B**) No suction feeding was observed in *nkx3.2*^-/-^ zebrafish (6 mpf); instead, they perform ram feeding (swim through food) with the fixed open gape. In this particular feeding episode, the mutant initiated a cycle with detection of food (change in swimming orientation) and turned laterally to exit that swimming trajectory in approximately 0.2 s. Frame by frame still images from a film captured at 60 frames per second. The movie is available as Supplementary Information (Movie S1).

With their modified skulls, adult *nkx3.2*^-/-^ mutants exhibited feeding behaviors markedly different from wildtype (Fig. 5; Movie S1). Wildtype zebrafish feed by suction using rapid lower jaw depression and a forward swing of the mobile premaxilla-maxilla complex shortly followed by opening of the operculum (Fig. 5A), consistent with general teleost feeding mechanics (Alexander, 1970, 1969; Lauder, 1980, 1979; Westneat, 2005, 2004). In contrast, adult *nkx3.2*^-/-^ mutants were constrained by the fixed upper jaw unit and open gape. They instead showed ram feeding behaviors (swimming through food) (Fig. 5B). The jaws remained fixed, and no significant dorsoventral movement was observed. Whereas in wildtype one complete cycle of the jaw opening and closing took approximately 80 milliseconds (and approximately a tenth of a second to the closure of the operculum), *nkx3.2*^-/-^ mutants required double that time from changing direction of swimming toward food (0 s) to doing so again away from the food (0.2 s) in this particular feeding episode in Fig. 5B. A detailed analysis of the feeding mechanics is beyond the scope of this paper and is currently the focus of our study, with filming at higher speed and resolution.

This ram-feeding behavior was correlated with skull remodeling in *nkx3.2*^-/-^ mutants. In zebrafish skulls, the bones form endochondrally (quadrate, anguloarticular, basihyal) or intramembranously (premaxilla, maxilla, dentary, jugal, opercular, preopercular) (Cubbage and Mabee, 1996; Schilling and Kimmel, 1997). Much of the skeletal remodeling observed in adult *nkx3.2*^-/-^ mutants occurred in the intramembranous bones — spatially and temporarily well outside the known expression domain of *nkx3.2* (Askary et al., 2017; Miller et al., 2003). Until past 1 mpf, the fusion between jaw cartilages was not completely ossified in these mutants, potentially allowing plastic remodeling (Fig. 2e, l). These observations suggest that *nkx3.2*^-/-^ zebrafish accommodate functional jawlessness through remodeling of the skull and changes to feeding behavior.

### *nkx3.2*^-/-^ zebrafish converge onto agnathans in overall head shapes

Through this dramatic remodeling of the skull, *nkx3.2*^-/-^ zebrafish assumed a head shape reminiscent of two lineages of extinct jawless vertebrates that have laterally compressed body profiles: 1) birkeniiform anaspids (Fig. 6C), stem cyclostomes known mostly from the Silurian period (Blom et al., 2001; Miyashita et al., 2019); and 2) furcacaudiid thelodonts (Fig. 6D), much more elusive stem gnathostomes known from the Silurian and Devonian periods (Märss et al., 2007; Wilson and Caldwell, 1998). Qualitatively, the resemblance is particularly striking in overall head shape characters, including: fixed gape (depressed lower lips), shortened snout, interorbital position of nostril, proportionally large branchial region, and massive parietal region behind the occiput.

**Fig. 6.**
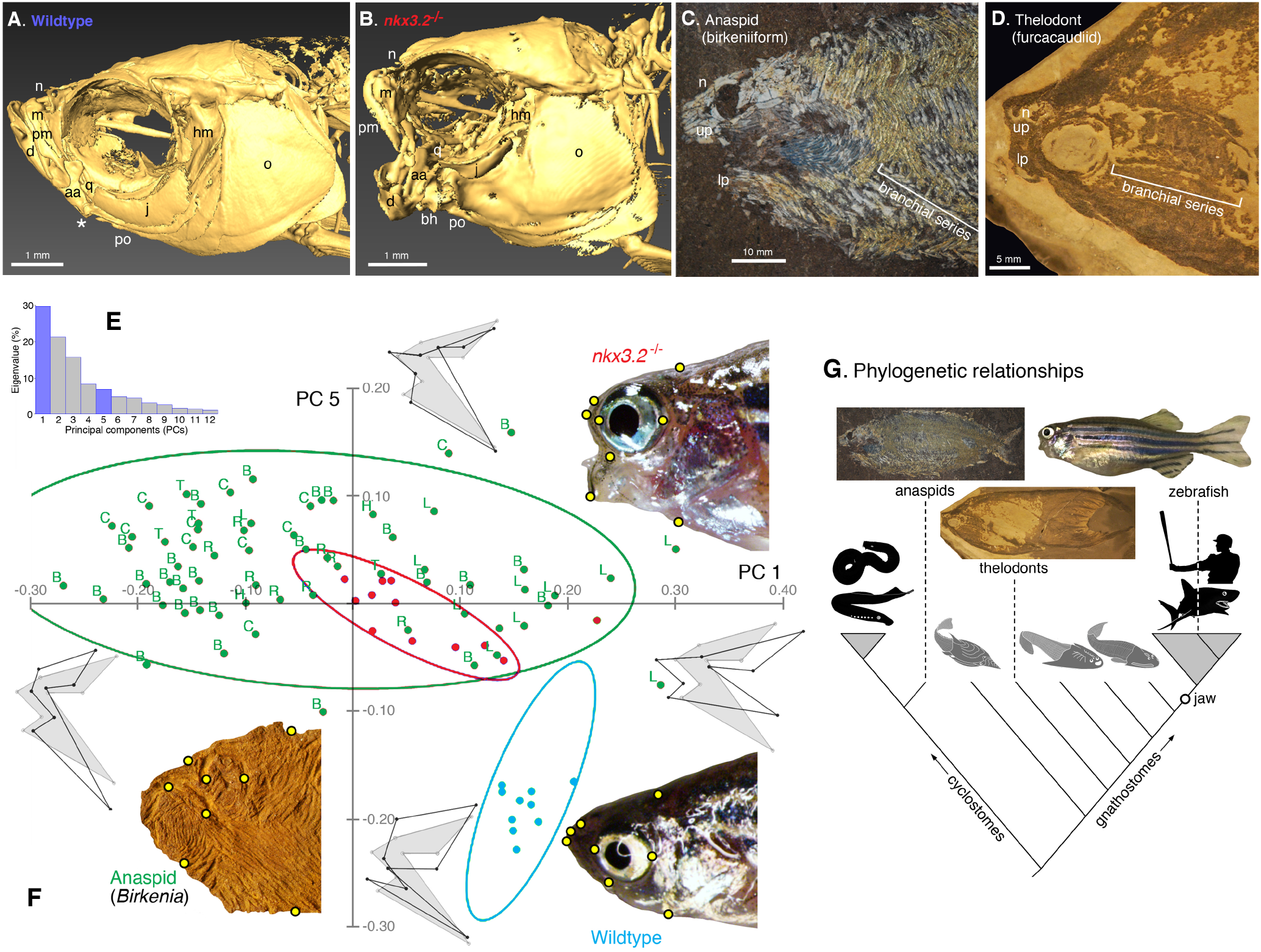
Adult *nkx3.2^-/-^* zebrafish converge onto overall head shapes of anaspids and thelodonts. (**A, B**) Skulls of adult zebrafish in left lateral view via micro-computed tomography (µCT) showing wildtype and *nkx3.2*^-/-^ null mutant zebrafish (allele *ua5011*; 20 bp deletion in homeodomain). The jaw joint is indicated by an asterisk (*) in wildtype (A) and absent in mutant (B). Adult mutants display dramatic phenotypes in the jaw, snout, lips, and orobranchial regions. See also 3D rendering in multiple angles (Movie S1). (**C**, **D**) Skulls of extinct jawless vertebrates, showing general resemblance to the skull shape of *nkx3.2*^-/-^ zebrafish. Morphometric analysis corroborates this similarity. (C) Skull of an anaspid (birkeniid birkeniiform) from the Upper Silurian Cape Philips Formation of Cornwallis Island, Canada (Geological Survey of Canada C-26661-005) in left lateral view. (D) Skull of *Sphenonectris turnerae*, a thelodont (furcacaudiid furcacaudiform) from the Lower Devonian Road River Formation of Northwest Territories, Canada (University of Alberta Laboratory for Vertebrate Palaeontology, specimen number 42212). (**E**, **F**) Landmark-based geometric morphometric comparison of *nkx3.2* phenotype using the thin-plate-spline Procrustes superimposition and principal component (PC) analysis. (**E**) Histogram showing loading on each principal component in eigenvalue (%). PCs 1 and 5 are highlighted in blue. These two PCs were chosen for comparison between groups, as these are the two largest components that set apart *nkx3.2* homozygous mutants and wildtype zebrafish from each other at adult stage. PCs 2-4, though accounting for greater variation than PC5, primarily distinguish among anaspids, or within wildtype and mutant zebrafish groups. The original dataset is available (Data Supplement 3). (**F**) A Cartesian plot of PCs 1 and 5, comparing morphospace occupation between wildtypes (2 mpf) (blue), *nkx3.2* homozygous mutants (2 mpf) (red), and anaspids (green), each with 90% ellipse. End of each axis is labeled with a thin-plate-spline shape at that position (dark outline) against mean shape (grey silhouette). A representative specimen is shown with landmarks labeled (yellow circles) for each group in left lateral view. Each data point is a unique biological replicate (a specimen). The anaspid example is *Birkenia elegans* (National Museum of Scotland specimen number 1929.5.6 from a Silurian locality of Scotland). For anaspids, each data point is labeled with taxonomic identifications: B, *Birkenia elegans*; C, Birkeniidae indet. from the Cornwallis Island; H, *Pharyngolepis oblongus*; L, *Lasanius problematicus*; R, *Ryncholepis parvulus*; T, *Pterygolepis nitidus*. (**G**) Simplified phylogenetic tree of vertebrates to illustrate distant relationships among the taxa compared in this paper. A 60 dpf gravid female specimen is shown for *nkx3.2*^-/-^ zebrafish (Fig. 3M). Photographs of an anaspid and a thelodont are the same individuals in the respective panels showing cranial morphology (D, E). Grey triangle indicates a crown group (not to scale). Dark silhouettes and long branches indicate crown groups, whereas grey silhouettes and short branches represent extinct lineages.

In linear morphometrics, the gape angle between upper and lower lips reveals that *nkx3.2*^-/-^ mutants closely resembles the anaspid- and thelodont-like conditions late in ontogeny (Fig. 4A). The trait in *nkx3.2*^-/-^ mutants departed from that in wildtype after the onset of skull ossification and active feeding (14–21 dpf), and overlaps with the range occupied by anaspids and thelodonts in later stages (1 and 2 mpf). When the jaws were at rest, the gape angle in wildtype zebrafish was consistently around 30 degrees regardless of ontogenetic stages. This was also the case for the *nkx3.2*^-/-^ mutants at 4 and 14 dpf when they still rely on yolk as main or partial intake. The gape angle increased steadily thereafter in the mutants.

In landmark-based geometric morphometrics, *nkx3.2*^-/-^ mutants aligned more closely with anaspids than wildtype along the principal components (PCs) that explain the adult phenotype (PCs 1 and 5) (Fig. 6E, F). Nearly one third of the overall shape variation loaded on PC1 and primarily concerned anteroposterior length and dorsoventral height of the entire head. Compared to wildtype zebrafish, *nkx3.2*^-/-^ mutants and most anaspids had a much shorter snout (with the lips and the nostrils shifting posteriorly toward the eye) and dorsoventrally deeper lower lips. PC 5 explained 7.0 % of overall shape variation, and the traits that vary along it were orientation of lower lip and relative height of nostrils, which clearly set wildtype zebrafish apart from *nkx3.2*^-/-^ mutants and anaspids. On the Cartesian grid of PCs 1 and 5, *nkx3.2*^-/-^ mutants overlapped with anaspids in morphospace occupation and apart from wildtype zebrafish (Fig. 6F). The area of overlap also indicated that *nkx3.2*^-/-^ mutants are broadly similar to anaspids in these PCs, not just to one or a few individual anaspid taxa. PCs 2-4 (not shown in Fig. 6) largely explain variation among anaspids or within wildtype/mutant zebrafish samples and were therefore uninformative for comparison between the groups (datasets containing landmark coordinates and Procrustes transformation are provided in Data Supplement 3).

## DISCUSSION

### Mutants corroborate the function of *nkx3.2* in joint development

Our *nkx3.2*^-/-^ zebrafish reinforce the morphant-based insight that this transcription factor is essential to the development of jaw joint in non-mammalian vertebrates (Miller et al., 2003). This is likely the widespread condition among jawed vertebrates. *Nkx3.2* knockdown also results in fusion between the palatoquadrate and Meckel’s cartilage in amphibians (Lukas and Olsson, 2018a), and a similar phenotype is observed in chicks with ectopic expression of BMP4 and FGF8 that repressed *Nkx3.2* (Wilson and Tucker, 2004). In mammals, the palatoquadrate (malleus) and Meckel’s cartilage (incus) migrate to form middle ear ossicles. The malleus-incus joint is not affected in *Nkx3.2*^-/-^ mice, even though the malleus becomes narrower in the mutants, and even though *Nkx3.2* plays a role in specification of the gonium and the anterior tympanic ring (Tucker et al., 2004). Aside from the head, *Nkx3.2* plays a role in various structures in mice and chicks, including the axial column (Herbrand et al., 2002; Lettice et al., 2001; Murtaugh et al., 2001) and visceral lateralities of the spleen and pancreas (Hecksher-Sørensen et al., 2004; Schneider et al., 1999). We will test in a forthcoming work whether or not *nkx3.2*^-/-^ zebrafish have parallel phenotypes to these amniote mutants in the axial skeletons or visceral literalities.

During development of the jaw joint, *nkx3.2* is thought to specify the joint interzone by inhibiting maturation or hypertrophy of the chondrocytes (Miller et al., 2003; Smeeton et al., 2016). Similar functions have been ascribed to *irx7* and *irx5a* in the hyoid joint (Askary et al., 2015). This is consistent with the predicted regulatory function of *Nkx3.2* in vertebral development, intervertebral and interphalangeal joint formation, or somatic chondrogenesis in general (Herbrand et al., 2002; Lettice et al., 2001; Murtaugh et al., 2001) — via repression of *Runx2*, by upregulating *Col2α1*, and/or through a positive feedback loop with *Sox9* (Smeeton et al., 2016). These insights are based on: a) experimental results using amniote embryos or somitic mesodermal cell cultures (Cairns et al., 2008; Kawato et al., 2012; Lengner et al., 2005; Murtaugh et al., 2001; Provot et al., 2006; Yamashita et al., 2009; Zeng et al., 2002); and b) clinical and genetic studies of human pathologies, including osteoarthritis and spondylo-megaepiphyseal-metaphyseal dysplasia (including pseudoepiphyses) (Caron et al., 2015; Hellemans et al., 2009). Before this study, however, no mutants were available to specifically address these potential mechanisms of *nkx3.2* functions in the mandibular arch.

### The open-mouth phenotype results from plastic remodeling

The late onset and topology of skull/jaw remodeling suggests that *nkx3.2*^-/-^ zebrafish accommodate the loss of the jaw joint via a plastic response. Other than the absence of jaw joint, *nkx3.2*^-/-^ zebrafish appear normal until metamorphosis (14–21 dpf) (Fig. 1D, G). As the skull ossifies, however, the observed phenotype becomes increasingly prominent (Fig. 2B, D, G, J, M, N). Although the fused jaw cartilages clearly result from *nkx3.2* mutation, *nkx3.2* loss-of-function, by itself, seems unlikely to yield all the rest of phenotypic effects described here. A genome-wide analysis suggests that *nkx3.2* patterning effects in the skull are restricted to the mid-portion of the mandibular arch (Askary et al., 2017). Nor does *nkx3.2* have known expression or function in the intramembranously ossified skull elements (Miller et al., 2003; Tucker et al., 2004), which are dramatically modified in our *nkx3.2^-/-^* fish. The hypertrophied mandibular cartilage may result from *nkx3.2* loss-of-function in regulating local chondrogenesis (Smeeton et al., 2016). Still, further investigation is warranted because in amphibians an ectopic cartilage formed in the mandibular arch with *Nkx3.2* overexpression, not repression (Lukas and Olsson, 2018b).

A comparative survey across vertebrates supports our interpretation of the adult *nkx3.2*^-/-^ phenotype as skeletal remodeling to accommodate the jaw joint loss. Discrete variation in cichlid jaw morphology has an epigenetic basis in behaviorally mediated skeletal remodeling, where gaping frequencies in juveniles correlate with dimensions of the retroarticular process (Hu and Albertson, 2017). The magnitude of morphological changes in the adult *nkx3.2*^-/-^ phenotype also appears consistent with a series of surgical experiments in mammalian jaw skeletons (Bayram et al., 2010; Gomes et al., 2012; Horowitz and Shapiro, 1955; Lifshitz, 1976; Miyazaki et al., 2016; Rodrigues et al., 2009; Sarnat, 1970; Sarnat and Muchnic, 1971; Toledo et al., 2014) or with the ‘bird face’ deformity observed in clinical cases of the temporomandibular joint ankylosis in humans (El-Sheikh et al., 1996). Collectively, these studies show that latent potentials of development allow a plastic trait to become expressed in jaw skeletons.

Variation resulting from developmental plasticities — whether induced by environmental cues, developmental perturbation, or mutation — are often non-random and adaptive (Palmer, 2012; West-Eberhard, 2005a, 2005b, 2003). Such non-random, adaptive responses are documented across wide ranges of taxa and structures, including: more robust claws in crabs fed with harder food items (Smith and Palmer, 1994); longer or shorter appendages of barnacles transplanted between wave-exposed and protected shores (Kaji and Palmer, 2017; Neufeld and Palmer, 2008); and thickened shells of gastropods exposed to predator cues (Appleton and Palmer, 1988; Edgell and Neufeld, 2008). Phylogenetically closer to zebrafish, cichlids have been extensively studied for developmental plasticity in jaw skeletons, such as: the relationship with gape frequencies and retroarticular processes, mentioned above (Hu and Albertson, 2017); antisymmetric development of the jaws in scale-eating specialists (Stewart and Albertson, 2010); and various dietary effects on feeding morphology (Galis, 1993; Greenwood, 1965; Liem and Osse, 1975; Meyer, 1987; Wimberger, 1992, 1991). Ram feeding facilitated by the fixed open gape in the adult *nkx3.2*^-/-^ zebrafish corroborates non-random, adaptive accommodation of the jaw joint defect.

### Jaw joint functions constrain skull morphology in vertebrates

The drastically remodeled skull of adult *nkx3.2*^-/-^ zebrafish highlights jaw movements as an important factor in the development — and thus morphological diversity — of jawed vertebrate skulls (Depew et al., 2005; Depew and Compagnucci, 2008; Depew and Simpson, 2006). Functional jaw loss resulting from *nkx3.2*-null mutations allowed the mutants to depart so markedly in morphology from their wildtype cousins, despite the nearly identical genetic backgrounds. This departure implies that movements at the jaw joint limit skull forms. The specific combination of traits observed in adult *nkx3.2*^-/-^ zebrafish — e.g., nostrils in interorbital position, premaxilla and maxilla abutted against antorbital wall, kinethmoid reduced, basihyal protrusion — is likely maladaptive and unavailable to zebrafish when the jaws function properly. Simultaneously, the absence of a jaw joint (or jaw apparatus altogether) also limits functionally viable forms. This interpretation is bolstered by the superficial convergence in head shapes between *nkx3.2*^-/-^ zebrafish and two Paleozoic agnathan lineages (anaspids and furcacaudiform thelodonts). The *nkx3.2*^-/-^ zebrafish and these agnathans share a functional requirement — the absence of a hinge joint between upper and lower lips — and transversely compressed body profile. No evidence suggests any more similarities in the otherwise wildtype-like young mutants (Fig. 1D, G) until the lower jaw skeletons begin rotating posteroventrally post 14 dpf (Fig. 2B, D). Therefore, we interpret the anaspid/thelodont-like traits in the adult *nkx3.2*^-/-^ zebrafish not as recapitulations of conserved, genetically hard-wired early vertebrate development (atavism), but rather as parallel developmentally plastic responses to shared growth conditions that were experienced by early vertebrates in a zebrafish skull (convergence).

Thus, jaw-joint loss (*nkx3.2* loss-of-function phenotype) released *nkx3.2*^-/-^ zebrafish from developmental constraints to facilitate jaw movement, and exposed them to a different functional requirement: feeding without mobile jaws. A fixed open gape — accompanied by modification of the skull elements — is one morphological solution to the functional loss of the jaw joint, which is corroborated by ram feeding exhibited by *nkx3.2*^-/-^ mutants (Fig. 5B; Movie S1). The magnitude of modification documented post-hatching (Figs. 1–5) is a testament to significant functional optimization within the bounds of the gnathostome bauplan. Simultaneously, well-constrained occupancy by the *nkx3.2*^-/-^ in the PCA plot (Fig. 6F) implies that alternative morphological patterns are either functionally non-viable (e.g., fixed closed gape) or developmentally non-accessible (e.g., ectopic formation of a mouth). Thus, the *nkx3.2*^-/-^ mutants provide a rare case study. The results suggest that the vertebrate bauplan allows a limited repertoire of functionally viable morphological patterns, onto which forms may converge under a given functional constraint.

In support of our interpretation, remarkable convergences arise via developmental plasticity under similar functional requirements — even in the absence of homology in individual skeletal elements — between taxa widely separated chronologically and phylogenetically. Bichirs (*Polypterus*) routinely trained under terrestrial conditions modify the pectoral fin skeleton through plasticity and develop morphological conditions observed in stem tetrapods (Standen et al., 2014). In another, *vav2* and *waslb* mutant zebrafish develop limb-like long bones connected by joints within the pectoral fins (Hawkins et al., 2018, preprint). West-Eberhard (2005b) reviewed a goat born with congenital paralysis of forelimbs, which performed bipedal locomotion. This behavioral accommodation led to highly modified musculoskeletal anatomy of the axial column, pelvis, and hindlimbs (Slijper, 1942a, 1942b). As shown in *nkx3.2*^-/-^ zebrafish, these experimental manipulations and clinical reports illustrate that adaptive forms emerge from developmentally plastic responses to functional constraints — and therefore may converge onto phenotype independently exploited by a distant lineage — even after development has laid out lineage-specific patterns (such as homologies of individual bones).

### *nkx3.2* mutants as a unique model for skeletal development and disease

Mechanisms of developmentally plastic remodeling, whether at the level of whole-animal behavior or gene transcription, remain an elusive component of skeletal development that is challenging to test experimentally. Therefore, *nkx3.2*^-/-^ zebrafish provide a unique experimental system in which to test further the role of a joint in skeletal development. For example: a) What genetic mechanisms regulate the remodeling of intramembranous ossifications? b) What components of the remodeling process respond to jaw-joint dysfunction? and c) Does any feedback exists to coordinate remodeling between developmentally independent, but functionally connected units (e.g., premaxilla-maxilla complex and kinethmoid)? Such insights would begin to fill in knowledge gaps regarding skeletal and joint diseases including osteoarthritis and joint ankylosis.

The non-atavistic nature of the *nkx3.2*^-/-^ phenotype is potentially useful to reevaluate evolutionarily inspired interpretations of phenocopies in general. There are no evolutionary relationships in the similarities between *nkx3.2*^-/-^ zebrafish and anaspids or thelodonts. Anaspids are a stem cyclostome lineage (Miyashita et al., 2019) (Fig. 6G), and thus a poor surrogate for an ancestral state. The lineages between anaspids and thelodonts, or those between thelodonts and the gnathostome crown, do not share a similar combination of morphological traits, but instead are dorsoventrally depressed forms (Janvier, 1996; Miyashita, 2016; Miyashita et al., 2019) (Fig. 6G). Finally, head similarities between the mutants and the stem taxa do not extend beyond overall configuration. No skull elements are lost or replaced in *nkx3.2*^-/-^ zebrafish to achieve the anaspid/thelodont-like micromery (Blom et al., 2001; Janvier, 1996; Märss et al., 2007). Therefore, the adult *nkx3.2*^-/-^ phenotype clearly does not represent a reversal to an ancestral state. Although experimental phenocopies are often interpreted as atavistic (ancestral reversal), alternative interpretations are seldom tested (Smith and Schneider, 1998). Here, our *nkx3.2*^-/-^ zebrafish present developmental plasticity as a testable alternative to atavistic reversal to explain a phenocopy. By showing that a gnathostome can survive without a jaw, the anaspid/thelodont-like *nkx3.2*^-/-^ zebrafish also offer a comparative model to make inferences about the functional morphology of these long-extinct agnathans.

## Acknowledgements

We thank B.F. Eames (University of Saskatchewan) for facilitating our collaboration; M.E. Bronner (California Institute of Technology), S.J. Childs (University of Calgary), M.I. Coates, V.E. Prince, and M.W. Westneat (University of Chicago) for discussion; and members of the Allison and Prince labs for maintenance of *nkx3.2^ua5011^*.

## Author contributions

T.M. conceived project; T.M. and A.P.O. performed experiments resulting in *nkx3.2^ua5011^*; J.S. and

N.N. performed experiments resulting in *nkx3.2^el802^*; P.B. and B.G. provided µCT scanning and histological sampling; T.M. and P.B. conducted morphometric analyses; T.M., P.B., A.R.P., J.G.C., D.G., and W.T.A. analyzed data; T.M. wrote the paper with inputs from P.B., A.P.O., J.S., A.R.P., J.G.C., D.G., and W.T.A.

## Supplementary information

Sequence information, measurements, and landmark coordinates are available as data supplements to this paper.

**Movie S1. Feeding behaviour and 3D µCT scans of jawless zebrafish.** Phenotypic comparison of *nkx3.2*^-/-^ and wildtype zebrafish. Three-dimensional rendering of µCT scan of the age-matched *nkx3.2*^-/-^ (ua5011) and wildtype (AB strain) zebrafish at 2 mpf, each followed by filming of feeding behavior on brine shrimp played at 1/10 original speed.

## Data supplements

**Data supplement 1.** Sequence information for the *nkx3.2* allele ua5011.

**Data supplement 2.** Gape angles and other linear measurements in zebrafish; gape angles and orbit diameter in anaspids.

**Data supplement 3.** Thin-plate-spline landmark data in zebrafish (wildtype and *nkx3.2*^-/-^) and anaspids used for Procrustes transformation, which were subjected to principal component analysis.

